# Multiplexed PCR to measure *in situ* growth rates of uropathogenic *E. coli* during experimental urinary tract infection

**DOI:** 10.1101/2024.11.21.624689

**Authors:** Santosh Paudel, Geoffrey B. Severin, Ali Pirani, Melanie M. Pearson, Mark T. Anderson, Evan S. Snitkin, Harry L.T. Mobley

## Abstract

Measuring bacterial growth rates *in vitro* is routine, however, determining growth rates during infection within a host has been more challenging. Peak-to-trough ratio (PTR) is a technique for studying microbial growth dynamics, calculated using the ratio of replication origin (*ori*) copies to those of the terminus (*ter*), as originally defined by whole genome sequencing (WGS). WGS presents significant challenges in terms of expense and data analysis complexity due to the presence of host DNA in the samples. Here, we used multiplexed PCR with fluorescent probes to estimate bacterial growth rates based on the abundance of *ori*- and *ter*-adjacent loci, without the need for WGS. We establish the utility of this approach by comparing growth rates of the uropathogenic *Escherichia coli* (UPEC) strain HM86 by WGS (PTR) and qPCR to measure the equivalent *ori:ter* (O:T^PCR^). We found that PTR and O:T^PCR^ were highly correlated and that O:T^PCR^ reliably predicted growth rates calculated by conventional methods. O:T^PCR^ was then used to calculate the *in situ E. coli* growth rates in urine, bladder, and kidneys collected over the course of a week from a murine model of urinary tract infection (UTI). These analyses revealed that growth rate of UPEC strains gradually increased during the early stages of infection (0–6h), followed by a slow decrease in growth rates during later time points (1-7 days). This rapid and convenient method provides valuable insights into bacterial growth dynamics during infection and can be applied to other bacterial species in both animal models and clinical infections.

**Importance:** Accurately measuring bacterial growth rates in the host, which plays a crucial role in determining the success of pathogens in establishing infections, poses significant challenges. To address this, bacterial replication rate has been measured as a proxy for growth rate estimation. While whole genome sequencing (WGS) has been used for this purpose, it comes with drawbacks such as high costs and difficulties in analyzing bacterial sequences due to the overwhelming presence of host DNA. In this study, we validate a more accessible PCR-based approach compared to the established WGS method and confirmed the reliability of our PCR-based technique. We then applied it to measure the growth rate of *Escherichia coli* during experimental urinary tract infection in a mouse model. This study provides a cost-effective and efficient alternative to WGS for studying bacterial replication dynamics during infection, potentially offering new insights into pathogen behavior and host-microbe interactions.

## Introduction

Urinary tract infections (UTIs) are among the most prevalent bacterial infections, with *Escherichia coli* being the primary causative agent constituting approximately 75% of uncomplicated UTIs and 65% of complicated UTI cases (1). While the virulence factors and pathogenicity of uropathogenic *E. coli* (UPEC) are generally appreciated (1–3), the *in vivo* growth dynamics during UTIs remain less well understood. Bacterial replication rate is important to pathogenicity as it can influence evasion of the host response, colonization, and persistence (4). Understanding bacterial growth rates within different infection niches will also advance our knowledge of how each unique environment affects bacterial proliferation and the progression of infection and helps pinpoint specific factors responsible for promoting or inhibiting bacterial growth.

Recent studies have established the use of bacterial circular chromosome replication as a means to determine growth rates with the underlying principle that rapidly growing bacteria have multiple replication forks starting at the origin (*ori*), leading to greater abundance of DNA proximal to the *ori* than distal sequences located near the terminus (*ter*) whereas slow growing or stationary phase bacteria have a more equitable number of *ori* and *ter* loci (Fig. 1). The ratio of sequencing coverage between the *ori* and *ter*, obtained from whole genome sequencing (WGS), is referred to as the peak-to-trough ratio (PTR) and was first applied by Korem *et al*. in 2015 for the determination of bacterial growth rates in a metagenomic study (5). Subsequently, a number of studies have utilized the concept of PTR to measure bacterial growth rates under a variety of conditions using WGS (5–12), digital droplet PCR (13), and SYBR green dye-based quantitative real time PCR (qPCR) (14, 15). While WGS-based PTR has been widely applied and allows for the quantification of both *ori* and *ter* adjacent sequence abundance within the same sample, this method is expensive, requires computational resources and expertise, and generates an unnecessary amount of sequencing data if growth rate determinations are the sole objective. While techniques utilizing qPCR-based methods offer a more economical approach to specifically quantify *ori* and *ter* adjacent sequences, their reliance non-specific DNA-intercalating dyes necessitates that the quantification of *ori* and *ter* abundance in the same sample be measured independently from one another. In our study, we combine both the ease of in-house qPCR with the WGS benefit of simultaneous measurement of *ori* and *ter* abundance in the same sample reaction by multiplexing qPCR reactions using TaqMan probes, and designate this O:T^PCR^ .

**Fig. 1.**
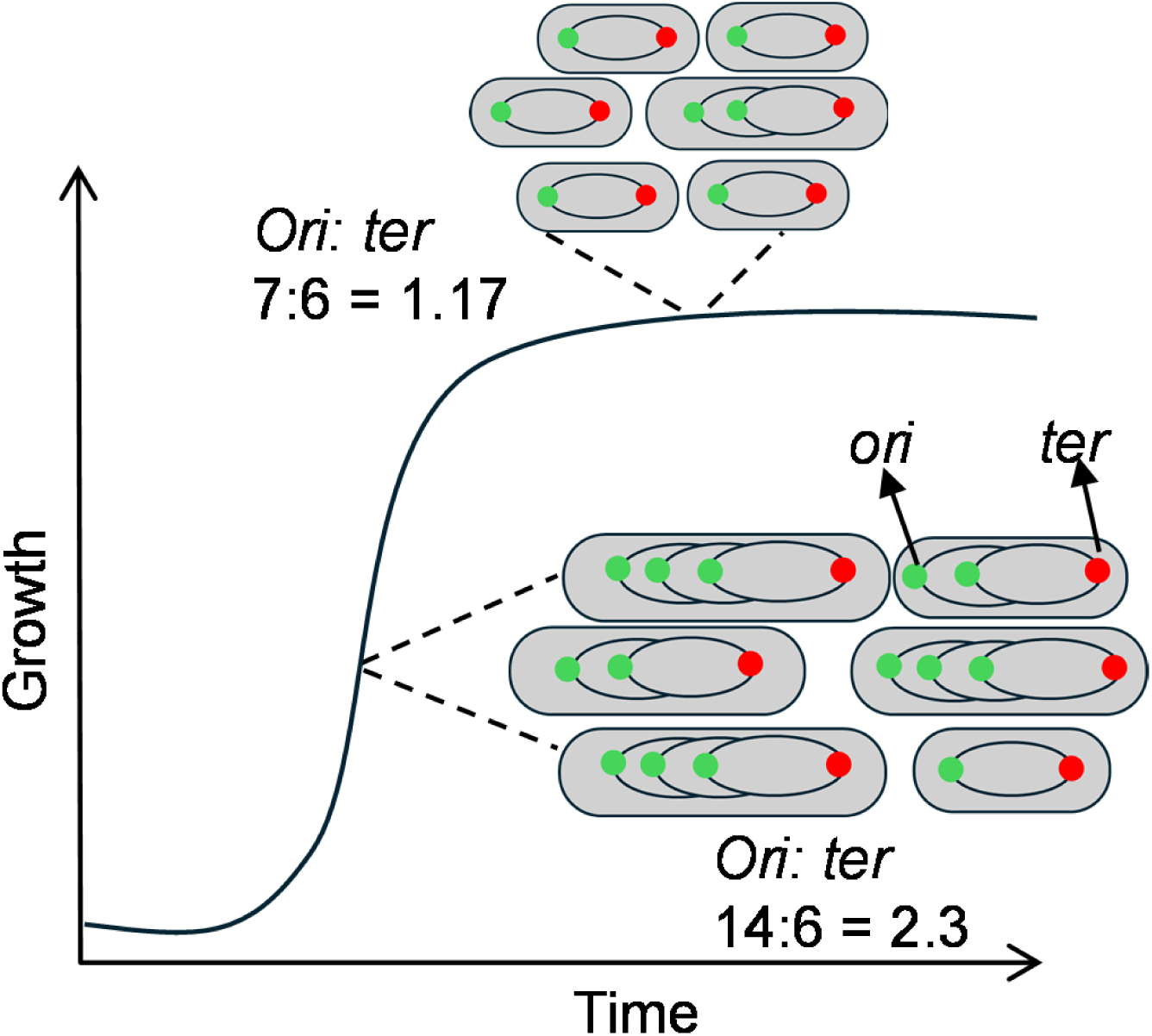
Determination of growth rate by PTR. Stages of bacterial growth representing their ori: ter (one of the examples of *ori: ter* during exponential and stationary phase is shown. *ori,* green*; ter,* red).

The purpose of this study was two-fold: 1) to develop a probe-based multiplexed qPCR method for measuring *ori:ter* and 2) to use this method to determine the growth rates of UPEC strains *in vivo* during experimental urinary tract infection in mice. Validation of this approach streamlines the process of determining *ori:ter* to accurately and rapidly estimates doubling times in diverse biological samples. Measuring bacterial growth rate *in vivo* within host tissue may provide insight into the dynamics of experimental infections over time and the possibly clinical outcomes. After validating our qPCR method with conventional PTR measured by WGS, we determined the growth dynamics of two UPEC strains HM56, and HM86 *in vivo* during experimental murine UTI for 7 days post inoculation. We found both UPEC strains rapidly replicating in murine urine sample early in the infection (6–12 hpi) with growth rates slowly decreasing through the course of a week. This study demonstrates the usefulness of applying a probe-based qPCR method to measure UPEC growth rates in a clinically relevant model of UTI, which could be leveraged for use in various types of infections and models.

## Materials and methods

### Bacterial strains and culture conditions

*E. coli* strains HM56 and HM86 were isolated from cases of uncomplicated UTI and have been described elsewhere (16, 17). Bacterial strains were revived from the frozen stock (-80°C) by culturing overnight in Lysogeny broth (LB), aerating at 200 rpm at 37°C. To conduct an *in vitro* experiment, the HM86 strain was cultured overnight with aeration at 37°C in LB, and inoculated 1:100 in fresh LB, terrific broth (TB), M9 minimal growth medium supplemented with 0.4% glucose (M9_Glc_), and pooled human urine (HU) from healthy female volunteers. We sampled bacteria from the culture at 0, 1, 1.5, 2, 2.5, 3, 3.5, 4, 5, and 6h to assess bacterial growth rate in different growth media, measuring absorbance (OD_600_) in cuvettes using GENESYS 10S VIS spectrophotometer (Thermo Scientific). We collected bacterial pellets for each time point by centrifugation at 17K x *g* (Fisher Accuspin Micro 17R) for genomic DNA. The bacterial pellet was immediately placed in dry ice and stored at -80°C until further processing. Bacterial growth rate was determined using the formula:

Growth rate (GR_p_) = (log(*OD*_p+t_/*OD*_p-t_))/dt, where “p” is the time point whose O:T^PCR^ values is correlated with the growth rate, “p-t” is the initial OD, “p+t” is the final OD, “dt” is the time elapsed between initial and final OD. For our growth rate calculation, t=30 minutes. Bacterial doubling time (DT) *in vitro* was calculated as [log(2)/GR_p_].

### Quantitative polymerase chain reaction (qPCR)

Genomic DNA (DNA) from *in vitro* bacterial culture samples and mice urine and organs were extracted using DNeasy® Blood and Tissue Kit (Qiagen, Germany) following manufacturer’s instruction. The qPCR was performed to determine the *ori:ter* ratio, a proxy of PTR (referred in this study as O:T^PCR^). The DNA concentration was estimated using Nanodrop One (Thermo scientific). The bacterial DNA in concentration of 500 pg was used to determine PTR for the *in vitro* samples. The targeted nucleotide sequences for the qPCR reaction near the origin of the replication (*ori*) is the *gidA* gene, and near the terminus of replication (*ter*) is the *dcp* gene.

The qPCR reaction was performed in QuantStudio 3 (Applied Biosystems) thermocycler (95°C 30 s, 40×(95 °C 5 s+58 °C 30 s). The optimized 25 µl reaction mixture contained 12.5 µl TaqMan^TM^ Fast Advanced Master Mix (Applied biosystems), 2.5 µl template DNA, and 10 µl of the reaction mixture of forward and reverse primer for *ori* and *ter* region (final primer concentration 1µM each), and probe for *ori* and *ter* sequence (concentration 0.25µM each). The annealing efficiency of the primer and probe binding to the *ori* and *ter* oligos was calculated using the equation [(E)= 10ၞ(-1/slope)-1] where E represents the annealing efficiency, and slope is derived from the standard curve of 10-fold dilution of DNA (5000 to 0.5 pg). To ensure the reliability and accuracy of the qPCR results, we repeated certain samples using a second set of primers that targeted the same region with the same probes for both set of primers. Primers (14, 18) and probes used for qPCR are provided in Table 1. The O:T^PCR^ was determined using 2^−ΔΔCt^ method (ΔCt= Ct_ori_ - Ct_ter_; ΔΔCt= ΔCt_test_- ΔCt_reference_). The PTR for each sample is the average PTR of 3 technical replicates. The sample of the same bacterial strain cultured to stationary phase of growth, whose peak-to-trough is expected to be 1, is used for the normalization during each run. To evaluate the specificity of the primers and probes utilized in the experiment we performed two experiments: 1) using DNA from various Gram-positive and Gram-negative bacteria, both individually and mixed with *E. coli* DNA (1:1) to determine whether the presence of DNA from other bacterial sources affect the *ori:ter* ratio, and 2) inclusion of DNA extracted from murine kidney to assess the *E. coli* DNA-specificity of the qPCR in the presence of mouse genetic material. Molecular biology grade water (Corning) was used as non-template control (NTC) for each cycling run. The sample tested for PTR using WGS was incorporated in every set of the qPCR performed to minimize the experimental variability in PTR measurements between experiments. The qPCR assay was performed for the UPEC strains cultured *in vitro* in LB, TB, M9_Glc_ and *in vivo* samples from the mouse during UTI.

**Table 1.**
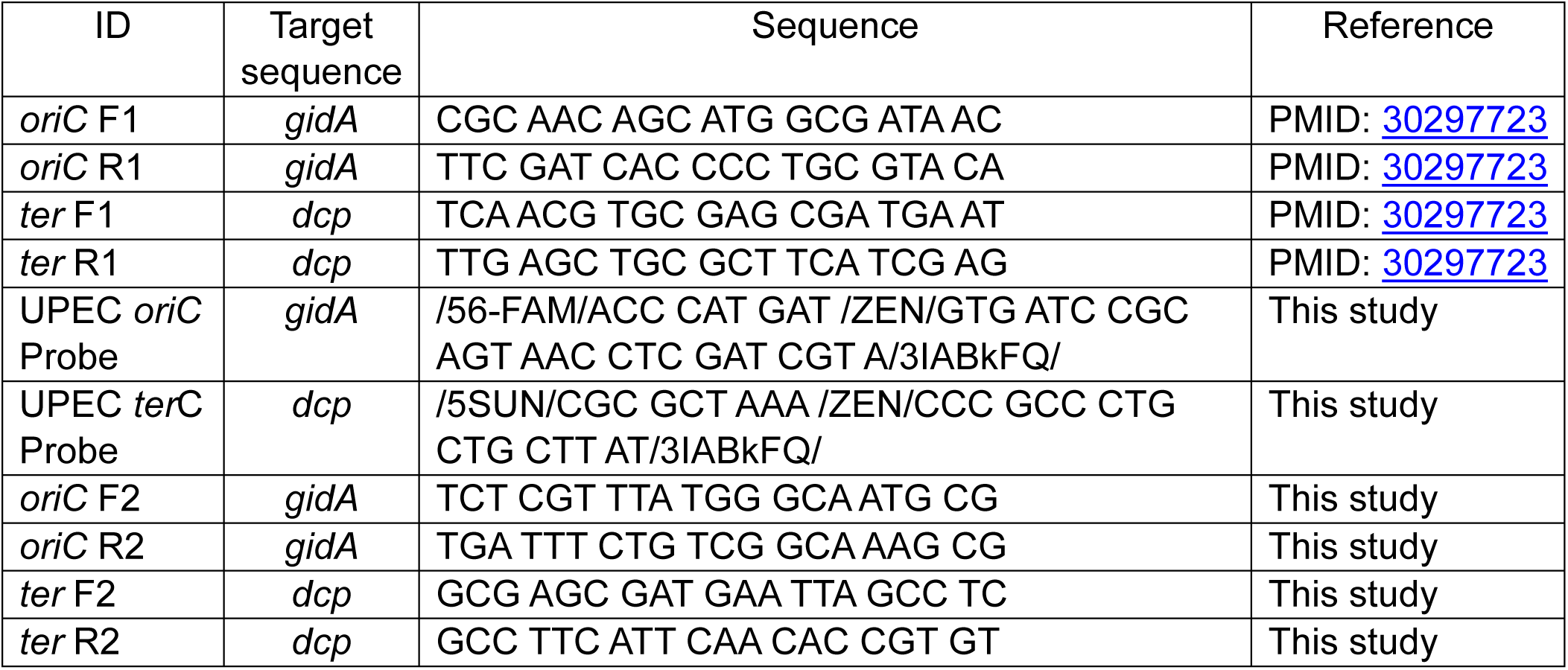
Primers and probes used in this study.

### Whole genome sequencing

PTR values were determined using previously published methods (5, 6). Briefly, sequenced reads from each sample were cleaned and mapped to the complete genome sequence of the originating bacterial isolate using Trimmomatic v0.39 and Bowtie2 v2.5.1. All alignments were indexed and sorted using SAMtools v1.18, and the coverage depth for each nucleotide position was extracted using Bedtools v2.30.0. A smoothing filter was applied to the mapped coverage of genomic segments, which was comprised of a moving sum with a window size of 10 kbp and a slide of 100 bp, followed by a moving median with a window size of 10-kbp bins and a slide of 100 bins. Instances where the bins did not have any mapped reads or had <50% remaining bins were discarded. The PTR was calculated from the peak and trough read coverage locations corresponding to maximum and minimum values, respectively, from the smoothed coverage (5). Snakemake implementation and analysis results can be accessed at https://github.com/alipirani88/Growth-rate-estimate_SMAKE. Assemblies for HM86 and HM68 were generated with Flye 2.9.2 and annotated with Prokka 1.14.5. Origin and termination of replication sites were determined by blasting qPCR OriC and Ter loci using BLAST 2.14.0

### Mouse model of ascending UTI

Six-to-eight weeks old female CBAJ (Jackson laboratory) mice were inoculated transurethrally with ∼2×10^8^ CFUs of UPEC strains as described previously (19, 20). Briefly, the overnight culture of UPEC strains cultured with shaking at 37°C was centrifuged for 30 minutes at 3000 rpm and the bacterial pellet was resuspended in sterile PBS. Mice were anaesthetized using ketamine/xylazine intraperitoneally (IP). Then, 50 µl of bacterial suspension was inoculated transurethrally using a sterile polyethene catheter connected to an infusion pump (Harvard Apparatus). After the induction of the UTI, urine samples from the inoculated mice were collected at 6, 24, 48, 72, 96, 120, 144, and 168 hpi for determination of CFUs and isolation of bacterial DNA. Mice were euthanized at 6, 24, and 168 hpi, and bladder and kidneys were collected for bacterial burden and bacterial DNA in the tissue samples. The bladder, and kidneys were collected in 5 ml culture tubes with 2 ml of ice-cold sterile PBS and homogenized using homogenizer (GLH 850, OMNI International). Bacterial burden from urine and each organ was enumerated using dilution plating by the drip-plate method and incubation at 37_°_C. The limit of detection (LOD) of bacteria in urine was 10^3^ CFU/ml, whereas for bladder and kidney is 20 CFU/organ. Urine and tissue homogenates were centrifuged at 17K x *g* at 4°C for 5 minutes. The supernatant was discarded, and the pellet was placed immediately on dry ice and stored at -80°C until further processing for DNA to measure bacterial growth rate determining PTR.

### Statistical analysis

Correlations between PTR and O:T^PCR^, and bacterial growth rate and O:T^PCR^ were evaluated using Pearson’s correlation coefficient. Linear regression analysis was performed to establish the relationship between growth rate and O:T^PCR^. Statistical significance of the doubling time (DT) measured by OD and PTR was determined using the paired *t*-test. To test the significance of *E. coli* burden in bladder, kidney, and urine as well as O:T^PCR^ in urine, we employed the Kruskal-Wallis test. Statistics and illustrations were performed using GraphPad Prism version 10 (GraphPad Software, CA, USA). A two-tailed *p*-value < .05 was considered statistically significant.

## Results

### Validating multiplexed quantitative real time PCR with whole genome sequencing as a tool for measuring origin to terminus ratios *in vitro*

To compare WGS PTR with our qPCR *ori:ter* method, we first generated a standard curve relating the growth rates of bacterial populations cultured *in vitro* to the relative abundance of the corresponding *ori* and *ter* sequences. This was achieved by culturing the *E. coli* strain HM86 in three different media where rapid and slow bacterial growth could be observed: lysogeny broth (LB), terrific broth (TB) and an M9 minimal medium containing glucose (M9_Glc_). Growth curves were initiated by addition of a one-to-one-hundred dilution of stationary phase HM86 culture into fresh medium and the optical density (OD_600_) of the bacterial population was measured at 30 to 60 minutes intervals over the six-hour period. Concurrent with these OD_600_ measurements, cell pellets were collected for genomic DNA extraction from which the relative abundance of the *ori* and *ter* sequences in the bulk population were measured by both multiplexed qPCR and WGS.

As anticipated, *E. coli* HM86 grew rapidly in both rich media (LB and TB) (Fig. 2 A) achieving exponential growth as early as 1.5 h post-inoculation and transitioning into stationary phase approximately two-hours later. In contrast, growth in M9_Glc_ was markedly slower with a longer lag phase and achieving a greatly reduced bacterial density over the course of six hours (Fig. 2A).

**Fig. 2.**
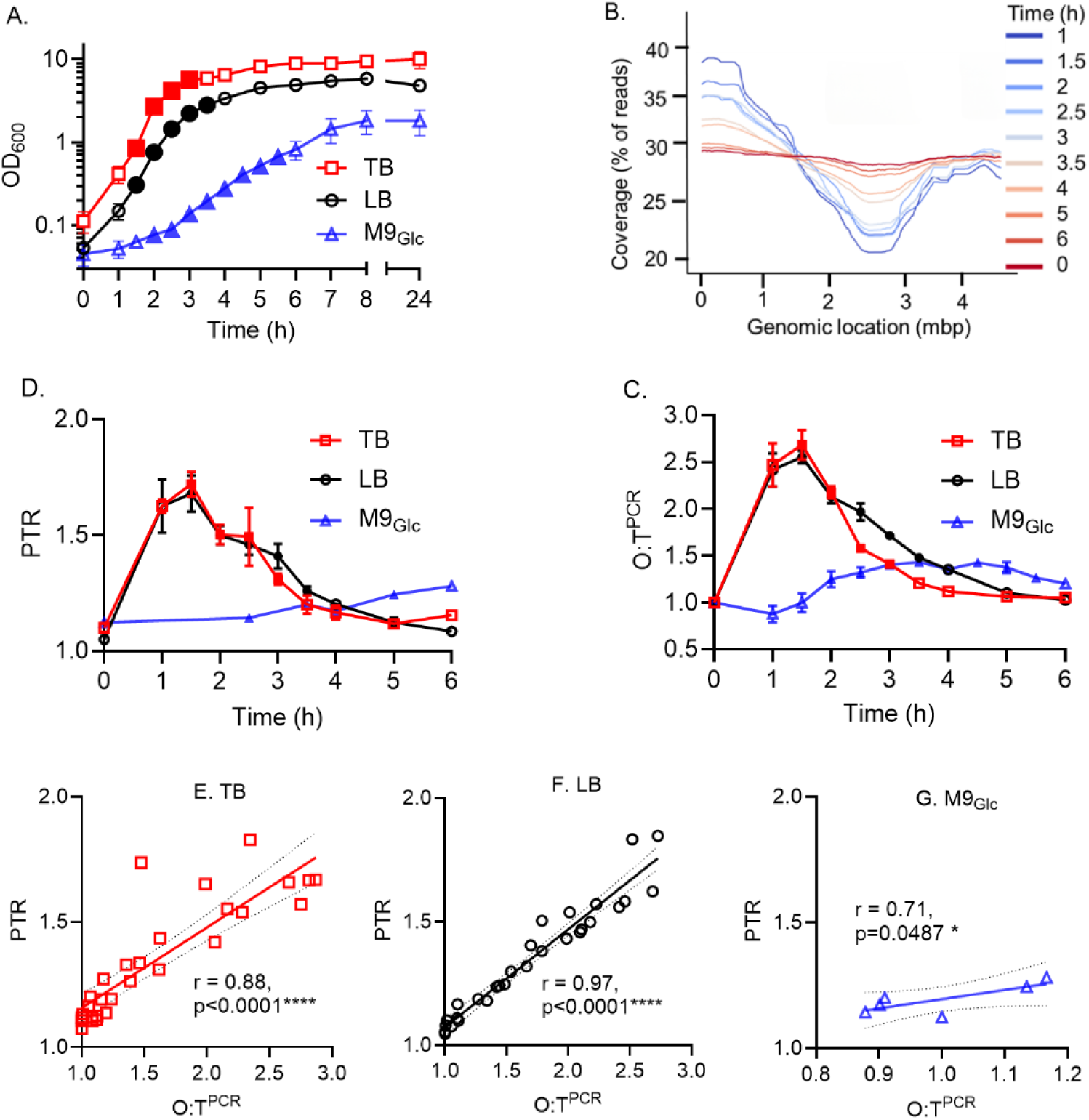
Validating multiplexed quantitative PCR with whole genome sequencing as a tool for measuring origin to terminus ratios in vitro. Growth experiments of *E. coli* strain HM86 in different growth medium conditions are plotted as OD_600_ reading (A) (mean ± SD). Means and standard deviation from three independent experiments are shown for TB (red), LB (black), and M9_Glc_ (blue). Three biological replicates for TB, and LB, and two biological replicates for M9_Glc_. (B) illustrates the example of the sequencing coverage across the genome of HM86 grown in vitro in LB up to 6h. Genomic location of origin (*ori*) and terminus (*ter*) of replication are denoted by peak and trough respectively. PTR determined by WGS is plotted in (C) (mean ± SD). (D) plots from the O:T^PCR^ values (mean ± SD). Scatterplot with linear regression for the relationship between PTR derived from WGS and *ori:ter* by qPCR method in TB (E), LB (F), and M9_Glc_ (G). N= 3 replicates (TB and LB), and N=1 (M9_Glc_). Pearson correlation coefficient (*r*) with *p*-values are shown in the graph. Statistical significance. **** p<.0001; *p<.05). Filled symbols in the graph A, C, and D indicates the data points with active replication included in the study for growth rate determination.

We first performed WGS on DNA samples collected throughout these growth curves to measure the relative abundance of *ori* and *ter* regions. Figure 2B illustrates one such read coverage plot from *E. coli* HM86 cultured in LB medium (Fig. 2A). During periods of rapid population expansion (1 – 3 h), we observed higher sequencing coverage (*i.e.*, peak) around the *ori* and lower sequencing coverage (*i.e.*, trough) corresponding with the location of the *ter*. As expansion of the population slowed, the relative abundance of reads mapping to the *ori* and *ter* regions normalized, resulting in a PTR of ∼1. Plotting the PTR from WGS data across the rich media growth conditions (Fig. 2C) illustrates the dynamic transition in PTR throughout the course of the growth curves (LB and TB, n = 3) with mean maximal PTR being observed at 1.5 h (LB = 1.68 ± 0.14, TB = 1.73 ± 0.1). While WGS was only performed for a single population of *E. coli* HM86 cultured in low nutrient M9_Glc_, the very modest changes in PTR predictably corresponded to the slow increase in culture density over time with a maximal PTR of 1.28 at 6 h (Fig. 2A).

We next tested whether multiplexing qPCR was a viable alternative to WGS for measuring the relative abundance of *ori* and *ter* sequences using these same *in vitro* DNA samples. Using previously published primers (14) and newly designed probes specific for regions near the *ori* and *ter,* we were able to rapidly generate *ori:ter* (O:T^PCR^) for *E. coli* HM86 cultured in all three media (Fig. 2D). Similar to PTR (Fig. 2B), the relative *ori:ter* measured by O:T^PCR^ in rich media were greatest during times of rapid population expansion with mean maximal *ori:ter* measured at 1.5 h (LB = 2.56 ± 0.12, TB = 2.69 ± 0.28) and this ratio gradually normalized to ∼1 as cultures reached stationary phase (Fig. 2A). We also observed that the *ori:ter* in M9_Glc_ also demonstrated a modest change in the O:T^PCR^ reflective of the slow and limited expansion of the *in vitro* population over time (Fig. 2A) with a mean maximal *ori:ter* of 1.43 ± 0.05 measured at 3.5 h. While the relationship between the *ori:ter* and *in vitro* culture growth phases were similar between PTR and O:T^PCR^ in all three media, the maximal values of the *ori:ter* during times of population expansion were greater when measured by multiplexed qPCR than WGS (Figs. 2C & 2D). Despite differences in the absolute values of O:T^PCR^ and PTR, we found the measurements *ori:ter* and PTR of each individual DNA sample were highly correlated in all three media conditions (Figs. 2E, 2F & 2G). We also performed the growth experiment of *E. coli* strain HM86 in human urine for 6h and measured PTR and O:T^PCR^ and found near perfect correlation of the measurements derived from WGS and qPCR methods (r=0.99, p=0.0002) (Fig. S3). The data revealed the slow growth of *E. coli* in vitro in human urine. The difference in absolute magnitude of *ori:ter* is likely reflective of a greater dynamic range in measuring *ori* and *ter* abundance by multiplexed qPCR and correlation of these measurements with PTR demonstrates the viability of using probe-based qPCR in lieu of WGS.

### O:T^PCR^ accurately predicts bacterial growth rate *in vitro*

Having demonstrated O:T^PCR^ is a reliable method to measure *ori:ter*, we then calculated growth rates for each time point across every growth curve based on the OD_600_ measurements that immediately preceded and followed that time point. A standard curve of growth rate and O:T^PCR^ was then established from all time points where the median growth rate from biological replicates cultured in same medium was less than 0.0025 (*i.e.*, doubling time of 120 min) (Fig. 3A). This minimum doubling time criterion functionally incorporated all regions of the three growth curves where active growth could be observed (*i.e.*, excluded lag and stationary phases) and the O:T^PCR^ were in excess of 1.25 and these time points are illustrated as filled symbols in Fig. 1A and 1D. From these data, we could construct a standard curve and derive a linear equation (Y=0.007648*X-0.006786) based on the relationship between growth rate and O:T^PCR^ (Fig. 3A). We then used this equation to calculate a predicted growth rate for all O:T^PCR^ measurements included in the standard curve and found no significant difference between the predicted growth rate and the experimentally derived growth from OD_600_ measurements for each O:T^PCR^ measurement (Fig. 3B). The constraints of this standard curve are the maximum and minimum O:T^PCR^ measurements of 2.87 and 1.16 representing a doubling times of 19.9 and 143.0 minutes, respectively. Therefore, O:T^PCR^ which are greater or less than these extremes cannot be used to accurately determine bacterial growth rates and are instead defined as extremely rapid growth or slow growth, respectively.

**Fig. 3:**
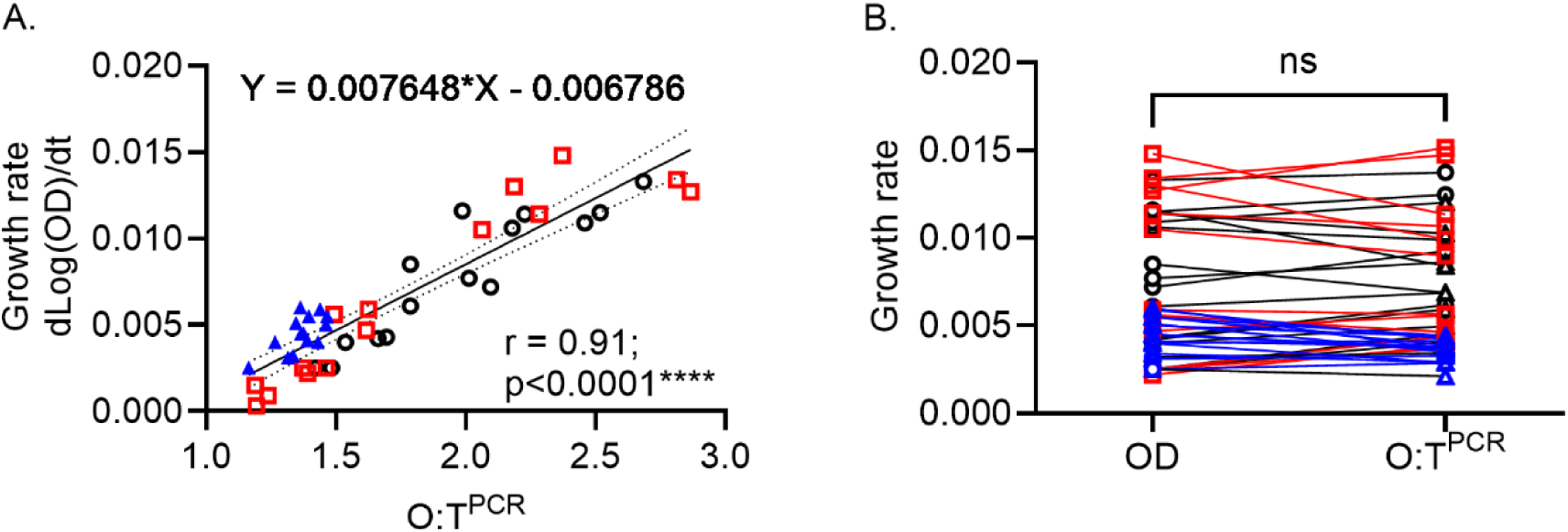
Bacterial growth rate correlates with O:T^PCR^. Graph represents the correlation between GR and *ori:ter*-values determined by qPCR method in TB+LB+M9_Glc_ (A). TB and LB=3 biological replicates, M9_Glc_= 2 biological replicates. Pearson correlation coefficient (*r*) with *p*-value is shown in the graph. Statistical significance: **** p<.0001; **p<.01). Bacterial growth rate calculated from the OD_600_ reading and the O:T^PCR^ in TB+LB+M9_Glc_ combined is plotted in Fig. B. Statistical significance was determined using the paired *t*-test, with *p*-value<0.05 considered a significant difference.

### Establishing limits of *ori:ter* quantification, detection, and specificity of O:T^PCR^

Under optimal conditions, such as bacterial monoculture, bacterial DNA can be extracted, quantified, and used in downstream molecular applications with the knowledge that all the nucleic acids present originated from a single source. In an infection model, DNA extracted from a complex biological sample will be a mixture of both host and etiological agent derived nucleic acids. Additionally, the concentration of bacterial DNA within this sample will be unknown and likely to be in low abundance. Therefore, we established two constraints for accurate *ori:ter* determination by O:T^PCR^ in complex biological samples: a limit of quantification (LOQ) establishing the minimum number of CFU (LOQ-CFU) from which DNA could be extracted and accurately measured, and LOQ-DNA demonstrating the minimum quantity of bacterial DNA which can be accurately measured.

Because the total number of CFU collected in a biological sample from an infection model can vary, we chose to establish LOQ which would represent the minimum number of bacterial cells required to provide adequate template DNA for accurate O:T^PCR^ determination. To establish this LOQ, we cultured *E. coli* HM86 to exponential phase in LB for 2 h from which 10-fold serial dilutions were prepared and DNA extractions of ∼10^6^, 10^5^, 10^4^, 10^3^, 10^2^, 10, and 0 CFUs were performed. From these DNA extractions we found the relationship between *ori* and *ter* was measured to be approximately 2.23±0.09 by O:T^PCR^ from templates originating from extractions containing 10^6^ to 10^3^ CFU, while those extracted from fewer CFU resulted in a significant deviation in this ratio (Fig. 4A). Therefore, an LOQ of ≥ 10^3^ CFU is required to provide sufficient bacterial DNA template for accurate determination of *ori:ter* by O:T^PCR^ .

**Fig. 4:**
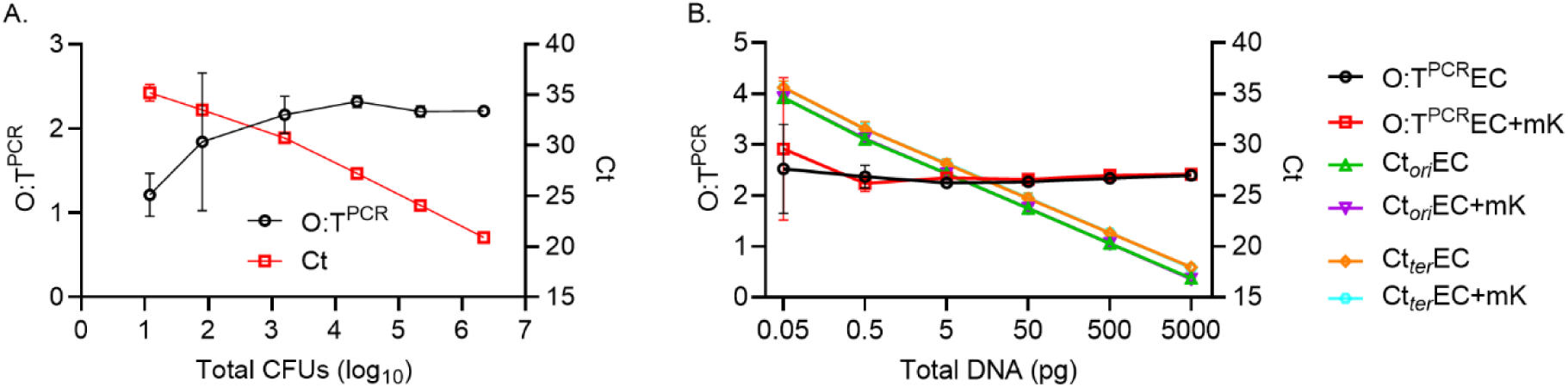
PCR assay optimization. Standard curves (Ct vs log_10_CFU) (A), and (Ct vs total DNA) (B) with their associated *ori:ter* determined by qPCR. Three replicates of the bacterial culture at exponential phase of growth (time point 2 h) was used for DNA extraction and determination of O:T^PC**R**^ [CFU vs *ori:ter* experiment (mean ± SEM)], and 3 independent experiment for total DNA vs *ori:ter* (mean ± SEM). The graph depicts *E. coli* strain HM86 *ori:ter*, Ct_ori_, and Ct_ter_ in presence and absence of murine kidney DNA (data from 3 biological replicate with 3 technical replicates) as indicated in figure legends. EC (*E. coli* DNA) and mK (murine kidney DNA). DNA (data from 3 biological replicates with 3 technical replicates) as indicated in figure legends. Ct (threshold cycle), EC (*E. coli* DNA) and kidney (murine kidney DNA).

In a biological sample collected from the infection model, bulk DNA extraction is likely to capture DNA from both the infectious agent and the host. This complex mixture of nucleic acids is likely to obscure the direct quantification of input template prior to its application in O:T^PCR^ . Because template abundance is correlated with critical threshold (Ct) values obtained by qPCR, we sought to define an LOQ based on the greatest Ct value for the *ori* specific oligo primers from which the relationship of *ori:ter* can be reliably determined by O:T^PCR^ . Using DNA extracted from ∼10^8^ CFUs, described in the LOQ experiment above, we performed O:T^PCR^ with 10-fold serial dilutions of template DNA (5 ng to 0.05 pg) to determine the range from which the *ori:ter* could be reliably measured. O:T^PCR^ containing 5 ng to 0.5 pg template DNA consistently measured an *ori:ter* of approximately 2.32±0.06, and this ratio deviated as the template abundance fell below this range (Fig. 4B). Over the entire range of templates tested, the *ori* oligo Ct values reliably increased by a mean Ct of 3.41 per 10-fold dilution of template (ideal change is 3.32 Ct per 10-fold dilution). The annealing efficiency of the primers and probes for *ori* and *ter* oligos were determined to be 96.2%, and 96.9 % respectively. Based on the these observations, we found the highest Ct value (*i.e*., lowest template abundance) from which *ori:ter* could be determined by O:T^PCR^ was 30.52±0.97 cycles. This LOQ establishes a critical threshold for informing the appropriate conditions for the application of *ori:ter* measured by O:T^PCR^; where the *ori* oligo Ct≤30.52, the corresponding *ori:ter* can be confidently used to accurately calculate the growth rate of the bacteria in that sample.

As described above, biological samples collected from an infection model are likely to contain at least two sources of DNA; the infectious agent and the host. There also exists the possibility that samples can contain DNA extracted from other sources (*e.g*., host commensals). To test the specificity of the *ori* and *ter* oligos to discriminate between *E. coli* and from other sources of DNA we performed O:T^PCR^ on mixed DNA templates containing 5 ng of *E. coli* HM86 DNA combined with of 5 ng of DNA isolated from alternative sources. In the absence of extraneous DNA, the template composed of exclusively *E. coli* HM86 DNA had a 16.04±0.13 *ori* oligo Ct and a 2.40±0.03 O:T^PCR^ . Addition of equal amount of DNA template purified from seven Gram-negative species cultured overnight (*Acinetobacter baumannii*, *Citrobacter freundii*, *Enterobacter hormaechei*, *Klebsiella pneumoniae*, *Morganella morganii*, *Pseudomonas aeruginosa*, *Serratia marcescens*), two Gram-positive species (*Bacillus subtilis* and *Staphylococcus aureus*) (Table 2), or host murine DNA (CBAJ) (Fig. 4B) did not significantly affect the detection of the *E. coli* HM86 template. While some reactivity with the *ori* and *ter* oligos was observed in the absence of *E. coli* HM86 DNA template for some of these alternative DNA samples, as well as the matrix control from the purification kit, the observed *ori* oligo Ct values were greater than the established LOQ above (≤30.5 Ct) (Table 2) and would therefore be excluded from further analysis.

**Table 2.** Specificity of *ori* and *ter* primers.

| **Template**<br>**DNA source** | **O:T<sup>PCR</sup>**<br>**(mean±SD)** | **Ct<sub>ori</sub>**<br>**(mean±SD)** | **Ct<sub>ter</sub>**<br>**(mean±SD)** |
| --- | --- | --- | --- |
| *E. coli* | 2.40±0.03 | 16.04±0.13 | 17.01±0.11 |
| *E. coli* + *Citrobacter freundii* | 2.44±0.07 | 16.08±0.12 | 17.07±0.08 |
| *E. coli* + *Klebsiella pneumoniae* | 2.61±0.16 | 15.94±0.09 | 17.03±0.04 |
| *E. coli* + *Serratia marcescens* | 2.57±0.04 | 16.00±0.03 | 17.08±0.01 |
| *E. coli* + *Proteus mirabilis* | 2.35±0.05 | 16.18±0.01 | 17.12±0.05 |
| *E. coli* + *Acinetobacter baumannii* | 2.57±0.02 | 15.89±0.16 | 16.96±0.14 |
| *E. coli* + *Enterobacter hormaechei* | 2.47±0.09 | 16.08±0.03 | 17.09±0.07 |
| *E. coli* + *Morganella morganii* | 2.46±0.03 | 16.01±0.07 | 17.02±0.06 |
| *E. coli* + *Pseudomonas aeruginosa* | 2.43±0.07 | 16.09±0.04 | 17.08±0.08 |
| *E. coli* + *Bacillus subtilis* | 2.41±0.03 | 16.09±0.12 | 17.07±0.14 |
| *E. coli* + *Staphylococcus aureus* | 2.46±0.19 | 15.93±0.18 | 16.93±0.08 |
| Matrix control | 0.98±0.06 | 34.53±0.03 | 35.95±0.37 |
| Naive mouse kidney | ND | 37.81±0.42 | >40 |
| *Citrobacter freundii* | ND | 35.5±0.36 | >40 |
| *Klebsiella pneumoniae* | ND | 30.24±0.07 | >40 |
| *Serratia marcescens* | ND | >40 | >40 |
| *Proteus mirabilis* | 1.43±0.44 | 35.64±0.47 | 37.32±0.93 |
| *Acinetobacter baumannii* | ND | 37.41±0.37 | >40 |
| *Enterobacter hormaechei* | ND | 37.2±0.55 | >40 |
| *Morganella morganii* | ND | 36.94±0.1 | >40 |
| *Pseudomonas aeruginosa* | ND | >40 | 39.47±0.04 |
| *Bacillus subtilis* | ND | >40 | >40 |
| *Staphylococcus aureus* | ND | >40 | 38.18±0.63 |

Note: ND=Not determined

**Table 3.** *E. coli* doubling time in murine urine during experimental UTI.

| Hours post-infection<br>(hpi) | Doubling time (minutes)<br>[Median (Min-Max)] |  |
| --- | --- | --- |
|  | HM56 | HM86 |
| 6 | 47.4 (31.6-105.2) | 54.2 (31.3-90.5) |
| 24 | 64.2 (28.9-153.1) | 67.6 (38.3-142.5) |
| 48 | 65.5 (36.8-70.9) | 70.4 (46.9-89.1) |
| 72 | 68.9 (59.8-148.0) | 63.6 (42.0-99.7) |
| 96 | 68.6 (55.3-128.1) | 73.1 (64.5-76.8) |
| 120 | 60.1 (46.4-70.1) | 56.3 (40.8-80.5) |
| 144 | 65.8 (58.5-68.4) | 86.1 (52.8-104.1) |
| 168 | 90.9 (63-129.1) | 75.8 (46.5-139.2) |

From these standardization and specificity experiments (Fig. 4), we found that *ori* and *ter* oligos are specific for *E. coli* HM86, extraction of >10^3^ bacterial CFUs is needed to provide enough template for O:T^PCR^, and a Ct≤30.5 for the *ori* oligos is required to accurately quantify the *ori:ter* in a given sample. By applying these criteria as quality control measures for evaluating bacterial growth rates we can apply O:T^PCR^ to complex biological samples collected from an infection model.

To ensure the reliability of our PCR method in measuring *ori:ter* ratios, we validated the same primer oligos with the other *E. coli* strain, HM56. We cultured strain HM56 in LB medium for 6h and measured O:T^PCR^ and found similar results to HM86, where HM56 DNA behaves like HM86 DNA indicating that our qPCR method is robust and applicable across different *E. coli* strains (Fig. S1).

### Mouse model of ascending UTI

Following parameterization of O:T^PCR^, we applied this method to measure *in vivo* bacterial growth rates in our model of interest, the experimental murine model of ascending UTI, over a period of seven days. For this, we induced experimental UTI in 6–7 week CBAJ female mice by transurethral catheterization and mono-culture inoculation of UPEC strains HM86 or HM56 isolated from uncomplicated female UTI patients (17, 21). Urine samples from the infected mice were collected at 6, 24, 48, 72, 96, 120, 144, and 168 hpi, whereas bladder and kidney samples were collected at 6, 24, and 168 hpi in a separate group of mice for bacterial burden and for extracting DNA and measuring growth rates from the two UPEC strains (Fig. 5).

**Fig. 5:**
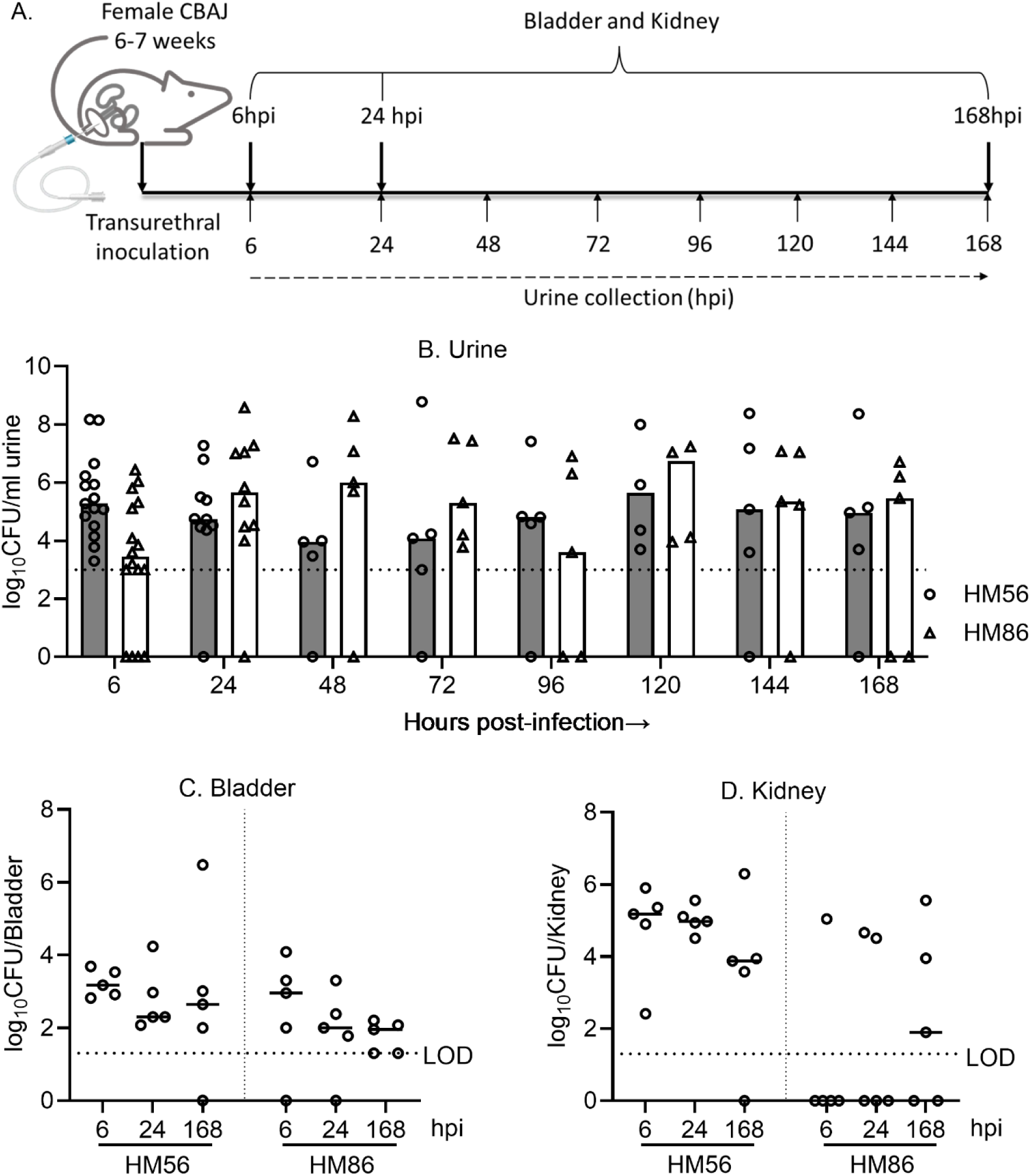
Bacterial burden in murine model of ascending UTI. Experimental plan for mouse model of ascending UTI (A). CBAJ mice (n=5 per time point) were inoculated with either HM56 or HM86 with ∼2*10^8^ CFU. Viable bacterial load in urine (B), bladder (C) and kidneys (D) were determined by dilution plating in LB agar at specific time point as mentioned in the plot. The horizontal dashed line indicates the limit of detection (LOD). Differences between time points were assessed by the Kruskal-Wallis test. Statistical significance differences are denoted by * where p<0.05.

Urine bacterial burden showed distinct patterns of colonization for each strain in the bladder, kidney and urine but no statistical significance difference in the CFU burden was observed over time. *E. coli* HM56 urine samples exhibited higher initial counts at 6hpi, with high abundance of bacterial culture later up to 7 days period. In contrast, *E. coli* HM86 exhibited lower urine bacterial count at 6hpi, increased by 24hpi, and maintained a constant level of persistence throughout the infection period (Fig. 5B).

Bladder colonization was similar for both strains, with higher level of colonization at 6hpi, which decreased by 24hpi, and remained at similar level up to 168hpi (Fig. 5C). The kidney colonization differed between the two strains. High level of kidney colonization was observed for *E. coli* HM56 at 6hpi, which was found to be lowered by approximately 10-fold at 168hpi. Conversely, *E. coli* HM86 exhibited limited kidney colonization early during the infection (6 hpi, 1 of 5 mice) which increased in abundance at later time points (24 hpi 2 of 5 mice; 168 hpi 3 of 5 mice) (Fig 5D) indicating a slower time to ascension. These findings highlight strain-specific variations in colonization patterns during the progression of UTI, providing insights into the dynamics of bacterial colonization, dissemination, and persistence in the different sites of the urinary tract.

### Growth rate of UPEC strains *in vivo* during UTI

Bacterial growth rates for HM86 and HM56 were next measured using DNA extracted from urine, bladder and kidney homogenates from infected animals. The recovery of bacterial DNA from the kidney sample was challenging due to the presence of host genetic material as evidenced by the observed Ct values compared to the recovered number of *E. coli* CFUs. However, the recovery was less affected in bladder and urine samples as observed Ct values were similar to the experiment performed with comparable known CFUs *in vitro* from pure bacterial culture. The DNA was extracted from urine, bladder, and kidney samples and *ori:ter* was determined. Fig. 6 illustrates the *ori:ter* values determined by qPCR from the urine samples during infection over the period of 168h. We observed comparable patterns of *ori:ter* between the two tested strains over the infection timeline, which shows the variability in the replication rate.

**Fig. 6:**
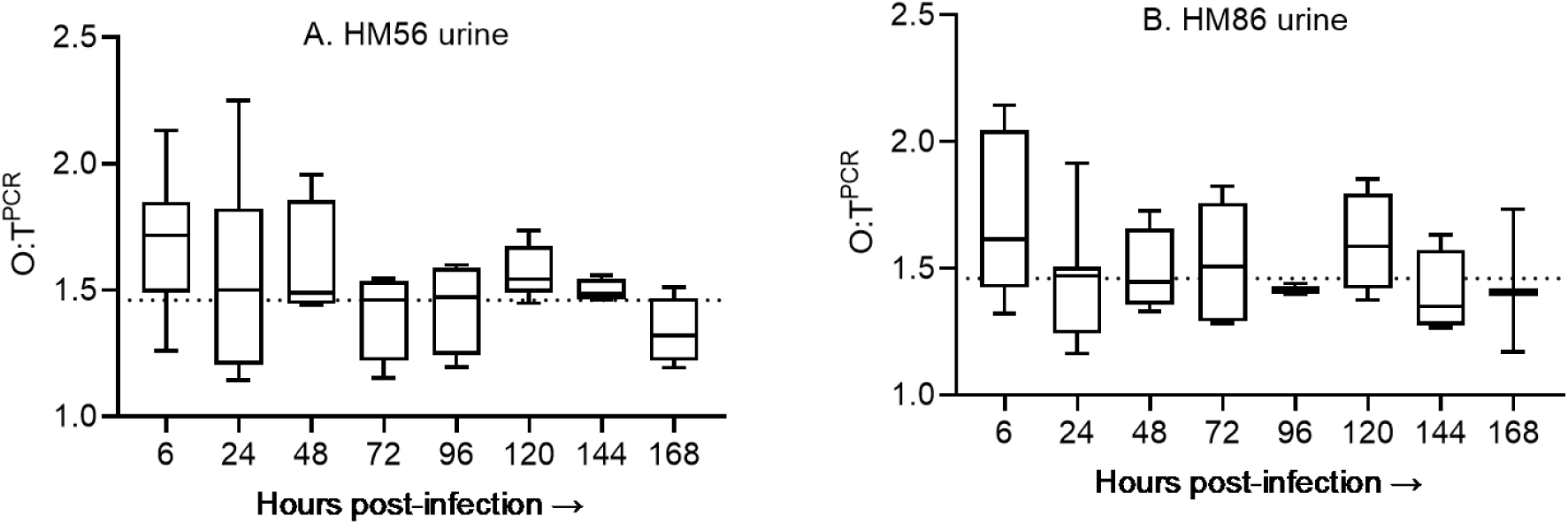
O:T^PCR^ in urine samples over infection period. Box and whiskers plots, where box represents the median and interquartile ranges and whiskers represents minimum and maximum values for each time during infection for HM56 (A), and HM86 (B). n= 5-15 mice per time point. Differences between time points were assessed by the Kruskal-Wallis test. Statistical significance differences are denoted by * where p<0.05.

From these *ori:ter* -values, we can predict that the growth rate of UPEC is higher at 6hpi, which gradually decreases over the subsequent infection period where the median *ori:ter* values is around our estimated borderline of the slow and fast growth conditions. No statistically significant difference was observed in *ori:ter* -values over the course of time.

The bacterial population doubling time was calculated from the qPCR derived *ori:ter*-values using the equation derived from the linear regression analysis of *E. coli* HM86 *in vitro* growth rate with their respective *ori:ter* value (Fig. 4D). The calculated *E. coli* DT in mouse urine is presented in Table 2. These DT results revealed a gradual increase in the growth rate of bacteria during the early stages of infection, peaking at 6h, followed by a slow decrease during later days transitioning to slower state of growth at 168hpi.

The fastest median DT for *E. coli* HM56 and HM86 is 47.4 and 54.2 minutes, respectively, observed for both at 6 hpi. The fastest growth of *E. coli* HM56 was observed at 24 hpi with a DT of 28.9 minutes, while for *E. coli* HM86, it was found to be 31.3 minutes at 6hpi. In case of individual mice, at 6hpi the fastest growth was determined as 31.6, and 31.3 minutes for HM56, and HM86 respectively. During the later time points, the UPEC strains exhibited variable growth rates. This trend suggests that UPEC undergoes rapid multiplication in the urine during early stages of UTI, particularly within the first 6h, followed by a gradual slowdown in growth rate. The similarity in growth rate between the two strains implies a similar growth strategy among different UPEC isolates during infection. Due to the presence of lower number of CFUs in most of the bladder, we were only able to determine *ori:ter* in few bladder samples with Ct value within the range of LOQ. For HM86, the DT in bladder was found to be 33.3 minutes at 6hpi (1/5 mice), which was found slower at 24hpi with DT of 61.2 minutes (1/5 mice). No bladder had sufficient CFUs for *ori:ter* determination at 168hpi for HM86. For HM56, in contrast to HM86, we observed slower growth at 6hpi with DT of 63.1 and 71.2 minutes (2/5 mice), which became even slower at 24 and 168 hpi with DT of 76.8 (1/5 mice) and 81.2 (1/5 mice) minutes respectively (Fig.S2-C). In contrast to the bladder, the growth rate in kidneys was found to be slower at early stage of infection, i.e., 6h followed by faster growth at 24 and 168 hpi for both *E. coli* strains. For HM56, the fastest growth rate was determined with DT of 76.6, 40.2, and 43.7 minutes at 6, 24, and 168 hpi, respectively. Since the kidney colonization with HM86 was limited, we were only able to determine the growth rate of 1 kidney samples at 6 and 168 hpi, with DT of 227.9 and 51.9 minutes, respectively (Fig. S2-D). Overall, the results revealed the infection-site specific differences in the *E. coli* growth rate over the course of infection period.

## Discussion

In this study, we developed a PCR-based method as a reliable alternative to WGS for determining bacterial growth rate targeting the specific regions near the origin and terminus of replication. As a proof-of-concept, the qPCR method was conducted on *in vitro* samples of *E. coli* HM86 cultured in both minimal and rich growth media and validated the *ori:ter* with results obtained from the WGS method. The validation experiments were performed using bacterial DNA isolated from the cultures grown under controlled laboratory conditions. This method provides an estimate of the population level PTR from the samples where bacteria are at different stages in their growth cycle. A series of the control experiments were conducted to validate the growth rate measurements by *ori:ter* using qPCR method for bacterial replication rates in host tissues. We determined that >10^3^ bacterial CFUs in pure culture condition gives accurate *ori:ter* measurements with the Ct value of *ori* oligos around 30.5, equivalent to 0.5 pg of the DNA used for *ori:ter* measurements. Therefore, we consider Ct values of 30.5 or higher to be unreliable for determining *ori:ter*. This results ensures that the *ori:ter* measurements remains accurate even when dealing with low bacterial loads in host tissues during infection studies. Using this validated method, we measured the *E. coli ori:ter* during experimental UTI from urine, bladder, and kidney samples from mice. DNA extraction and *ori:ter* measurement in urine and bladder samples with the bacterial count above 10^3^ CFUs resulted Ct value within the range of cut-off Ct value providing reliable result. This might vary as the recovery of DNA varies between the extraction protocol. Below this threshold, the consistency of the *ori:ter* was compromised, rendering inconsistencies of the result. In terms of DNA concentration, a total DNA amount of 0.5 pg was sufficient to produce a consistent *ori:ter* value, demonstrating the method’s high sensitivity. We were only able to measure *ori:ter* from limited bladder samples since only a few bladder samples had sufficient bacterial counts to enable reliable *ori:ter* measurements within reliable range of Ct values. Although the colonized kidney samples had higher number of CFUs count than that required for accurate *ori:ter* measurement, we were not able to measure *ori:ter* from some kidney samples as the Ct values derived from the qPCR method were higher than the cut-off Ct values. This, however, was not due to the interference of the mouse kidney DNA to reading for bacterial DNA as evidenced by the experiment performed by adding 50 ng of the mouse kidney DNA to 0.5 pg of *E. coli* DNA without deviation in the Ct values and *ori:ter* measurements. This might be due to the inability of our DNA extraction protocol to recover enough bacterial DNA from the mouse kidney tissue homogenate.

Experiments using two sets of primers targeting the same region yielded similar results, confirming the reliability of the performed method. Since we employed a probe-based qPCR method, it was highly specific in amplifying the target sequence from the intended bacterial species. This approach allows us to amplify and measure two targeted chromosomal regions from a single sample using their specific probes, thereby reducing discrepancies in readings between wells for the replication origin and terminus regions when determining *ori:ter* measurements. This multiplexing capability is advantageous in reducing both costs and result discrepancies compared to running samples in separate wells for the two target regions.

From the *in vivo* growth rate experiment during UTI in mice, we observed active growth in the urine samples with *ori:ter* >1 in all samples. The fastest growth was observed during the early phase of infection for the tested UPEC strains which gradually slowed during later days. Previously, the growth rate *in vivo* was determined in mouse UTI as well as human patients by determining PTR, a sequencing-based method, which revealed that UPEC strains exhibit rapid growth during human UTIs, often matching or exceeding *in vitro* rates (6). Similar to this study, DT in mouse urine and bladder during experimental UTI at 6hpi was determined to be 34.9, and 36.9 minutes respectively, with much slower growth rate at 24hpi (6).To measure the growth rate *in vivo* during UTI, another study employed qPCR-based method targeting *ori* and *ter* region (22) from the urine sample of the infected human patients with UTI, where active growth with day-to-day and inter-patient variability in growth rate was observed (22). Notably, *E. coli* strains isolated from the urinary tract demonstrate significantly higher growth rates in urine compared to fecal isolates, suggesting adaptation to the urinary environment (6). The slow growth observed during the later stage of infection in the urine and bladder might be due to the multifaceted adaptive response to the challenging host environment such as activation of bacterial stress response mechanism, increased pressure from the host immune system, formation of protective biofilm, or the development of metabolically dormant persister cells. By slowing their growth, *E. coli* can manage nutrients to persist under nutrient-limited conditions, evade host immune responses, and maintain a viable population capable of resisting elimination and potentially causing recurrent infections. This strategy underscores the complex interplay between bacterial pathogens and host defenses in the context of chronic or recurring UTIs (23–29). In contrast to urine and bladder, the *E. coli* growth rate in the kidney was found to be higher during later stages of infection which highlights the niche-specific variations in growth pattern within different areas of the urinary system. The distinct growth dynamics in the kidney suggest that this organ provides a unique environment for *E. coli* proliferation as the infection progresses. From the bacterial burden and growth rate results, we did not observe the direct connection between growth rate and total number of CFUs recovered from urine, bladder, and kidneys.

The strategy of measuring *ori:ter* to study bacterial growth rate has been applied in other infection models, such as mouse peritonitis, revealing heterogeneous growth rates within bacterial populations during infection (14), and blood-stream infections (30) showing organ specific pattern of growth rate in multiple bacterial species. The method has also been tested with slower-growing pathogens like *Xylella fastidiosa*, showing promise for assessing growth status in plant and insect vector environments (31). Importantly, this approach has also been applied to predict antibiotic treatment efficacy (32).

In conclusion, our research establishes the effectiveness of a probe-based qPCR method for measuring *ori:ter* and *in situ* determinations of bacterial growth rate in a cost-effective and efficient manner. This approach of measuring bacterial growth rate by PCR has potential applications in various infection models across different bacterial species.

## Acknowledgements

We would like to acknowledge the members of the Mobley lab and Bachman Lab for their insightful feedback throughout this project. This work was supported by the funding NIAID R01AI165582 (H.L.T.M.).

## Notes

### Competing Interest Statement

The authors have declared no competing interest.

### Summary of Updates

Measuring bacterial growth rates in vitro is routine, however, determining growth rates during infection within a host has been more challenging. Peak-to-trough ratio (PTR) is a technique for studying microbial growth dynamics, calculated using the ratio of replication origin (ori) copies to those of the terminus (ter), as originally defined by whole genome sequencing (WGS). WGS presents significant challenges in terms of expense and data analysis complexity due to the presence of host DNA in the samples. Here, we used multiplexed PCR with fluorescent probes to estimate bacterial growth rates based on the abundance of ori- and ter-adjacent loci, without the need for WGS. We establish the utility of this approach by comparing growth rates of the uropathogenic Escherichia coli (UPEC) strain HM86 by WGS (PTR) and qPCR to measure the equivalent ori:ter (O:TPCR). We found that PTR and O:TPCR were highly correlated and that O:TPCR reliably predicted growth rates calculated by conventional methods. O:TPCR was then used to calculate the in situ E. coli growth rates in urine, bladder, and kidneys collected over the course of a week from a murine model of urinary tract infection (UTI). These analyses revealed that growth rate of UPEC strains gradually increased during the early stages of infection (0-6h), followed by a slow decrease in growth rates during later time points (1-7 days). This rapid and convenient method provides valuable insights intobacterial growth dynamics during infection and can be applied to other bacterial speciesin both animal models and clinical infections.

