## Supplemental Files for "Multiplexed PCR to measure *in situ* growth rates of uropathogenic *E. coli* during experimental urinary tract infection"

### Supplementary figures

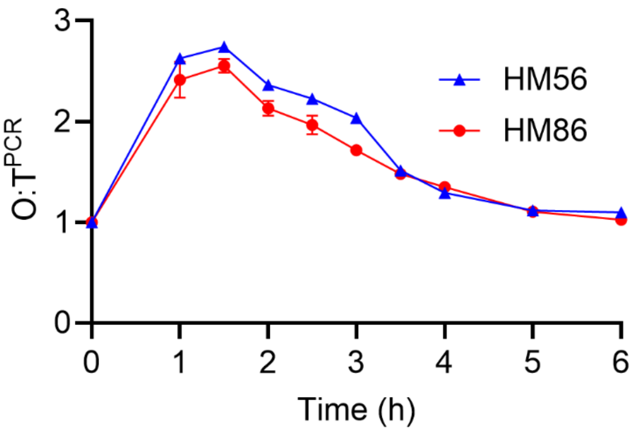


**Fig.S1. O:T$^{PCR}$ of the UPEC strains HM86 and HM56 cultured in LB medium**. (n=3 for HM86, and n=1 for HM56).

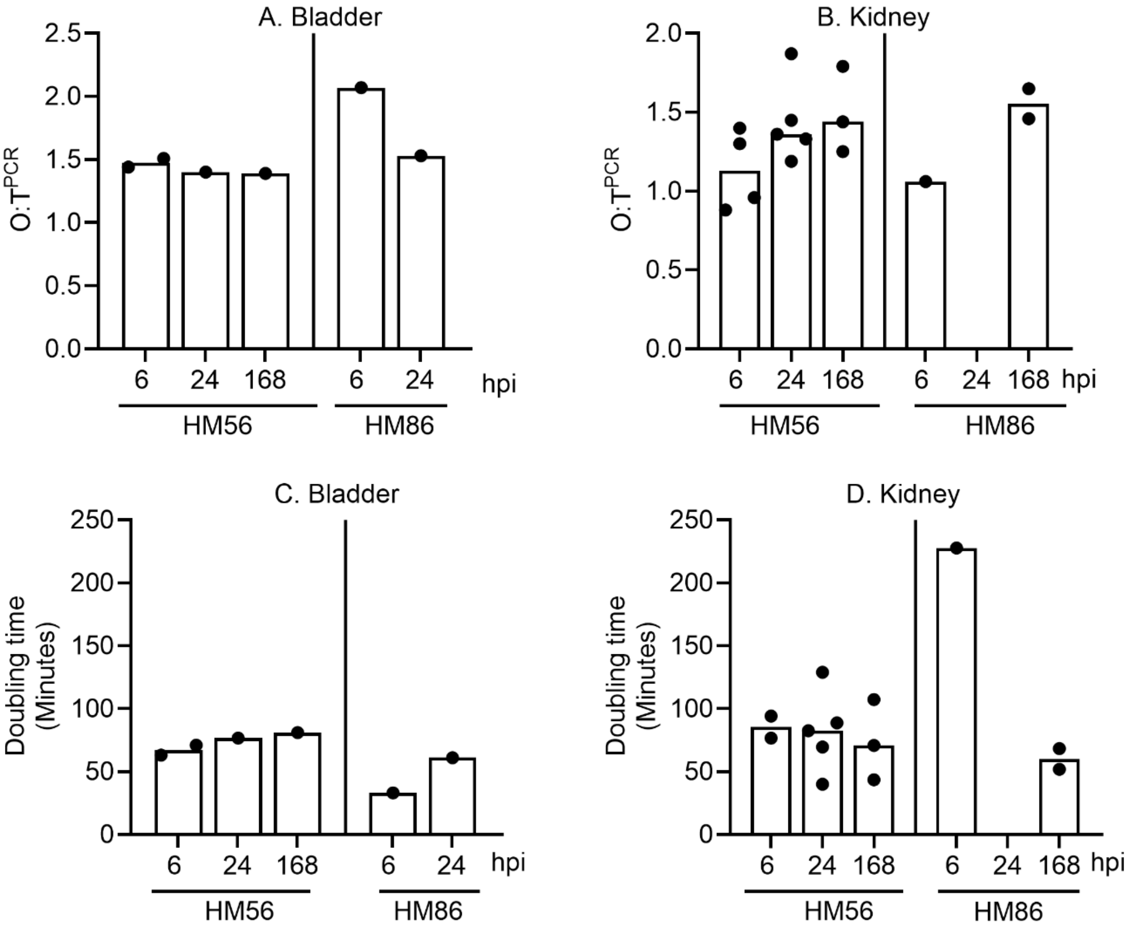


**Fig. S2. O:T$^{PCR}$ (Bladder-A, and Kidney-B) and *E. coli* doubling time (Bladder-C, and Kidney-D) of the mouse during experimental UTI at the time point of 6, 24, and 168 hpi.**

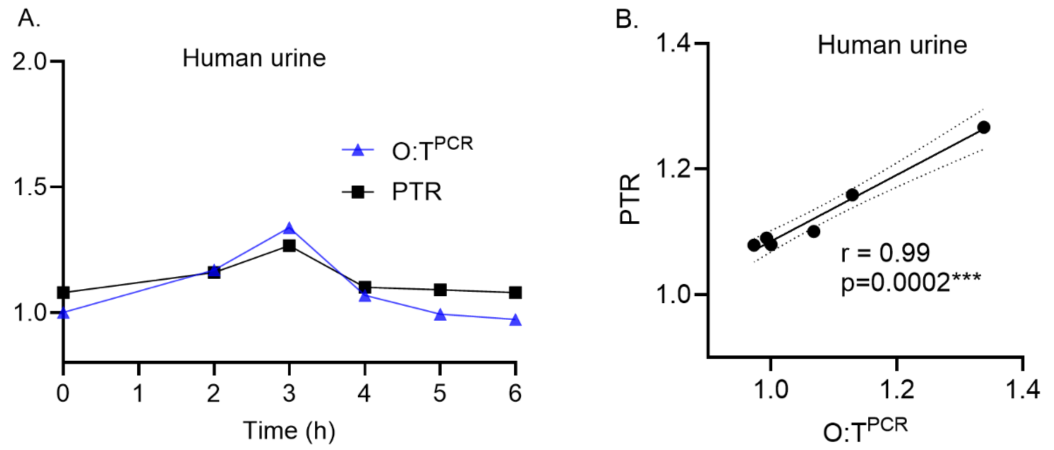


**Fig. S3. O:T$^{PCR}$ and PTR of E. coli strain HM86 cultured in human urine (A), and scatterplot with linear regression for the relationship between PTR derived from WGS and *ori:ter* by qPCR method (B).** Pearson correlation coefficient (*r*) with *p*-value is shown in the graph. Statistical significance. *** p<.001).
